# Development of a Deep Learning model Tailored for HER2 Detection in Breast Cancer to aid pathologists in interpreting HER2-Low cases

**DOI:** 10.1101/2024.07.01.601397

**Authors:** Pierre-Antoine Bannier, Glenn Broeckx, Loïc Herpin, Rémy Dubois, Lydwine Van Praet, Charles Maussion, Frederik Deman, Ellen Amonoo, Anca Mera, Jasmine Timbres, Cheryl Gillett, Elinor Sawyer, Patrycja Gazińska, Piotr Ziolkowski, Magali Lacroix-Triki, Roberto Salgado, Sheeba Irshad

**Author notes:** Corresponding author Pierre-Antoine Bannier. These authors jointly supervised this work: Roberto Salgado and Sheeba Irshad. Author Details: Glenn Broeckx, Loïc Herpin, Rémy Dubois, dwine Van Praet, Charles Maussion, Frederik Deman, Ellen Amonoo, Ancar Mera, Jasmine Timbres, Cheryl Gillett, Elinor Sawyer, Patrycja Gazinska, Piotr Ziolkowski, Magali Lacroix-Triki, Roberto Salgado, Sheeba Irshad.

## Abstract

**Introduction:** Over 50% of breast cancer cases are "Human epidermal growth factor receptor 2 (HER2) low breast cancer (BC)", characterized by HER2 immunohistochemistry (IHC) scores of 1+ or 2+ alongside no amplification on fluorescence in situ hybridization (FISH) testing. The development of new anti-HER2 antibody-drug conjugates (ADCs) for treating HER2-low breast cancers illustrates the importance of accurately assessing HER2 status, particularly HER2-low breast cancer. In this study, we evaluated the performance of a deep learning (DL) model for the assessment of HER2, including an assessment of the causes of discordances of HER2-Null between a pathologist and the DL model. We specifically focussed on aligning the DL model rules with the ASCO/CAP guidelines, including stained cells’ staining intensity and completeness of membrane staining.

**Methods:** We trained a DL model on a multi-centric cohort of breast cancer cases with HER2- immunohistochemistry scores (n=299). The model was validated on 2 independent multi- centric validation cohorts (n=369 and n=92), with all cases reviewed by 3 senior breast pathologists. All cases underwent a thorough review by three senior breast pathologists, with the ground truth determined by a majority consensus on the final HER2 score among the pathologists. In total, 760 breast cancer cases were utilized throughout the training and validation phases of the study.

**Results:** The model’s concordance with the ground truth (ICC = 0.77 [0.68 - 0.83]; Fisher P = 1.32e-10) is higher than the average agreement among the 3 senior pathologists (ICC = 0.45 [0.17 - 0.65]; Fisher P = 2e-3). In the two validation cohorts, the DL model identifies 95% [93%- 98%] and 97% [91% - 100%] of HER2-low and HER2-positive tumors respectively. Discordant results were characterized by morphological features such as extended fibrosis, a high number of tumor-infiltrating lymphocytes, and necrosis, whilst some artifacts such as non-specific background cytoplasmic stain in the cytoplasm of tumor cells also cause discrepancy.

**Conclusion:** Deep learning can support pathologists’ interpretation of difficult HER2-low cases. Morphological variables and some specific artifacts can cause discrepant HER2-scores between the pathologist and the DL Model.

## Introduction

Approximately 15-20% of breast cancer patients exhibit HER2 gene amplification or overexpression, resulting in aggressive tumor behavior and resistance to conventional therapies^1–3^. HER2-positive breast cancer, defined as IHC 3+ or IHC 2+ and in situ hybridization (ISH)+, is treated with specific HER2-targeted therapies. Conversely, HER2- negative breast cancer, defined as IHC 0, IHC 1+, or IHC 2+/ISH−, historically did not qualify for HER2-targeted therapy. However, some tumors traditionally categorized as HER2- negative have low levels of HER2 expression, termed HER2-low breast cancer, defined as IHC 1+ or IHC 2+/ISH−. Recently, DESTINY-Breast04 trial has shown that the HER2-targeted antibody-drug conjugate trastuzumab deruxtecan (T-DXd) improves survival in patients with previously treated advanced or metastatic HER2-low breast cancer^2^. Therefore, it is critical to assess HER2 status accurately in daily practice. HER2-low breast cancers are cases that do not fit neatly into the traditional positive or negative categories. These cases show no HER2 amplifications, yet they display detectable levels of HER2 expression on IHC (1+ or 2+/ISH-). These patients can benefit from trastuzumab deruxtecan^2^.

This new HER2-low paradigm has challenged the pathology community. The need for more accurate HER2 testing at the low end of the HER2 spectrum is of importance. Intratumoral heterogeneity of HER2-staining can make the pathologist’s assessment more difficult ^3,4^. Pathologists’ ability to accurately classify HER2 IHC, with minimal inter-observer variability, is paramount for selecting those patients for HER2-targeted therapy^5^.

In addition to the challenges posed by HER2-low breast cancer cases, recent technological advancements, particularly in artificial intelligence (AI) and digital image analysis (DIA), offer promising solutions. These technologies have the potential to assist pathologists in interpreting challenging HER2 cases with greater accuracy and efficiency. The co-use of AI- assisted assays alongside pathologists could mitigate both inter- and intra-pathologist variability, leading to more consistent and reliable HER2 status assessments^6^.

In this study, we propose the development of a deep-learning model that aligns methodologically with the ASCO/CAP criteria for HER2 assessment. Our analysis focuses on identifying morphological features and artifacts contributing to ambiguity in HER2-low assessment, aiming to enhance the precision of HER2 status determination in daily practice. By leveraging advanced computational techniques, we seek to provide pathologists with a robust tool for more accurate classification of HER2 expression, ultimately improving patient selection for HER2-targeted therapy.

## Results

Within the study, we included a total of 760 patients diagnosed with breast cancer. We built a deep learning model to predict the HER2 IHC score (0, 1+, 2+, 3+) from IHC slides. The model underwent training utilizing a cohort of 299 breast cancer cases (named Cohort 1) and subsequent validation across two external validation cohorts: Cohort 2 (n=369) and Cohort 3 (n=92). Out of the training set, 80 cases lacked HER2 status annotations and were consequently excluded from subsequent analysis. This resulted in a total of 219 cases available for training the model in Cohort 1. Given the inter-observer variability of HER2 assessment^6^, we ensured that in our validation cohorts, HER2 evaluations were conducted by three experienced pathologists specifically trained in assessing HER2 expression levels according to the 2018 ASCO-CAP guidelines (**Figure 1**). Consecutive IHC and H&E slides of 515 cases of breast carcinomas were collected and digitized from the validation cohorts (**Figure 1a**). Pathologists carried out a visual inspection to rule out pairs of slides with artifacts (pen markers, folded tissue, out-of-focus regions). Following this review, 59 (59/369) slides from Cohort 2 and 18 (18/92) from Cohort 3 were excluded due to significant slide artifacts (**Supplementary Table 1**). In Cohort 2, the majority of excluded cases were excluded due to “pen markers” on the slides, and in Cohort 3 due to low-quality staining. The average agreement between the 3 expert pathologists reached an intra-class correlation coefficient of 0.45 [0.17 - 0.65] (Fisher P = 0.002) for the IHC 0 and 1+ cases (**Figure 1b**).

**Figure 1:**
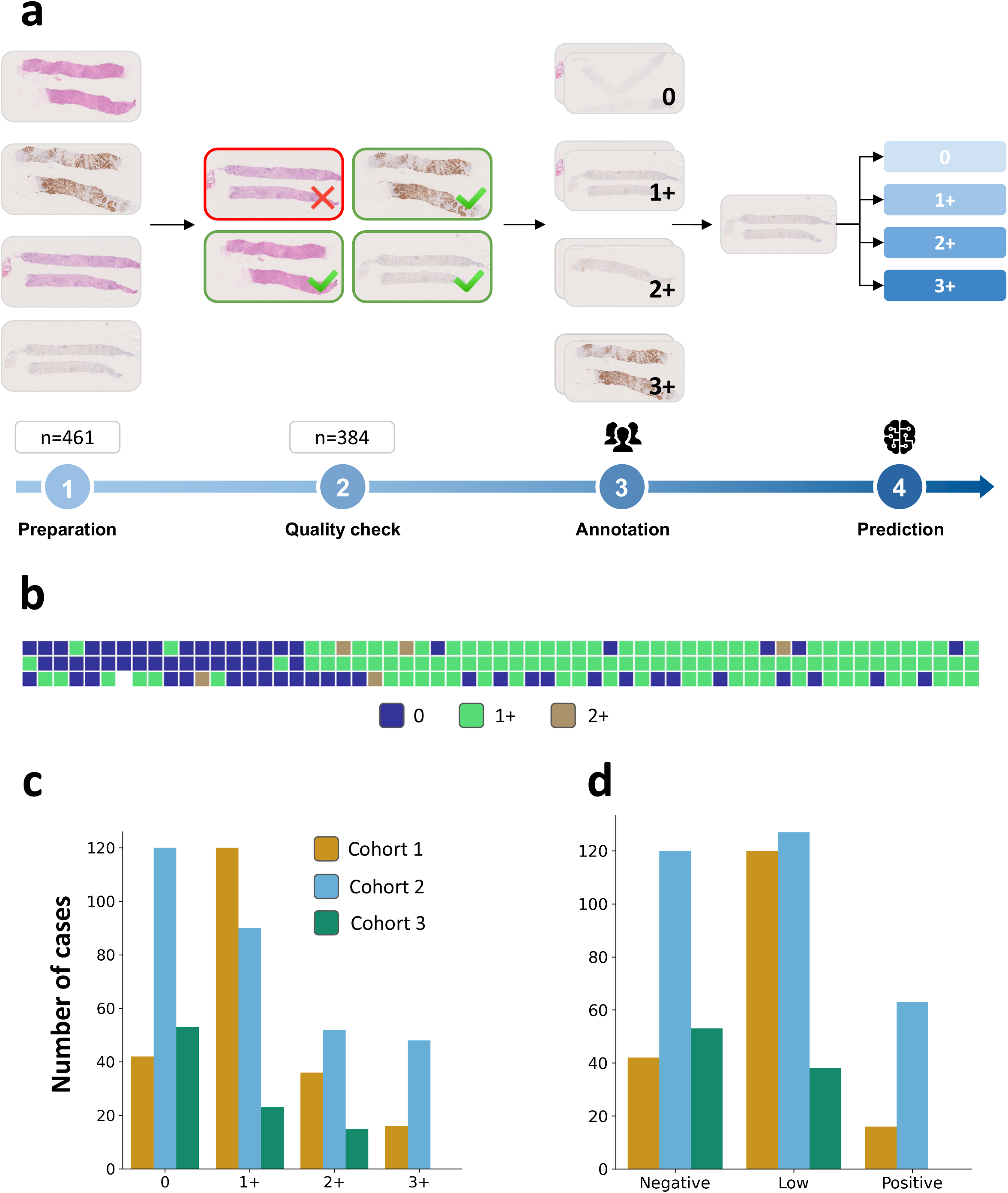
Clinical workflow and validation strategy. a) Consecutive IHC and H&E slides of 461 cases of breast carcinomas were collected and digitized in multiple centers (Cohort 2 and 3). A visual inspection was carried out by pathologists to rule out pairs of slides with artifacts (pen markers, folded tissue, out-of-focus regions). For the slides passing the quality check, 3 expert breast pathologists review the IHC slides and gave a score (0 / 1+ / 2+ / 3+) by applying the 2018 ASCO/CAP guidelines for HER2 quantification. The final IHC score was determined as the majority vote among the pathologists. For equivocal cases (IHC 2+), an ISH test was performed to verify ERBB2 amplification, allowing us to map equivocal cases as HER2+ or HER2-low. The HER2 status was determined via IHC and imputed to the corresponding H&E slides. A model was developed to quantify the IHC score from the IHC slides. **b)** Agreement plot between the 3 annotators for the validation cohort for 50 cases annotated either IHC 0 or IHC 1+. The 3 annotators reached an intraclass correlation coefficient of 0.45 [0.17 - 0.65] (p = 0.002) for the IHC 0 and IHC 1+ cases of Cohort 2. **c)** Distribution of the HER2 IHC score across the different cohorts. **d)** Distribution of the HER2 status across the cohorts.

Distribution of IHC scores and HER status for the 3 cohorts are shown in **Figures 1c** **& 1d**. The IHC scores across the three cohorts were: Cohort 1: 0 = 40/219 (18%), 1+ 120/219 (55%), 2+ 39/219 (18%), 3+ 19/219 (9%); Cohort 2: 0 = 120/369 (33%), 1+ 87/369 (24%), 2+ 53/369 (14%), 3+ 50/369 (13%), unknown (low quality) = 59/369 (16%); Cohort 3: 0 = 54/92 (59%), 1+ 22/92 (24%), 2+ 16/92 (17%), 3+ 0/92 (0%) (**Figure 1c**). The HER status across the three cohorts were: Cohort 1: Negative = 40/219 (18%), low 122/219 (55%), positive 19/219 (9%), unknown (no FISH) = 38/219 (18%); Cohort 2: Negative = 120/369 (33%), low 120/369 (33%), positive 66/369 (18%), unknown (low quality) = 59/369 (16%); Cohort 3: Negative = 54/92 (59%), low 38/92 (41%), positive 0/92 (0%) (**Figure 1d**). Particular attention was drawn not to include too many IHC 3+ cases, which are considered easily recognizable.

### The model’s concordance with the ground truth improves the average agreement among senior pathologists

Next, we trained our model to predict the IHC score (0, 1+, 2+, or 3+) on Cohort 1 (n=219). This model successfully determined the HER2 protein on two independent validation cohorts (AUC: Cohort 2 = 0.95 [0.93 - 0.96], Cohort 3 = 0.92 [0.90 - 0.93]; **Figure 2a, 2b**; **Supplementary Table 2**. The model reached an intra-class correlation coefficient of 0.95 [0.94 - 0.96] (Fisher P = 1.32e-10) on the entire cohort and 0.77 [0.68 - 0.83] (Fisher P = 5.43e- 7) on cases annotated as 0 or 1+, improving the average agreement between the 3 senior pathologists. The confusion matrix is presented in **Supplementary Table 3**.

**Figure 2:**
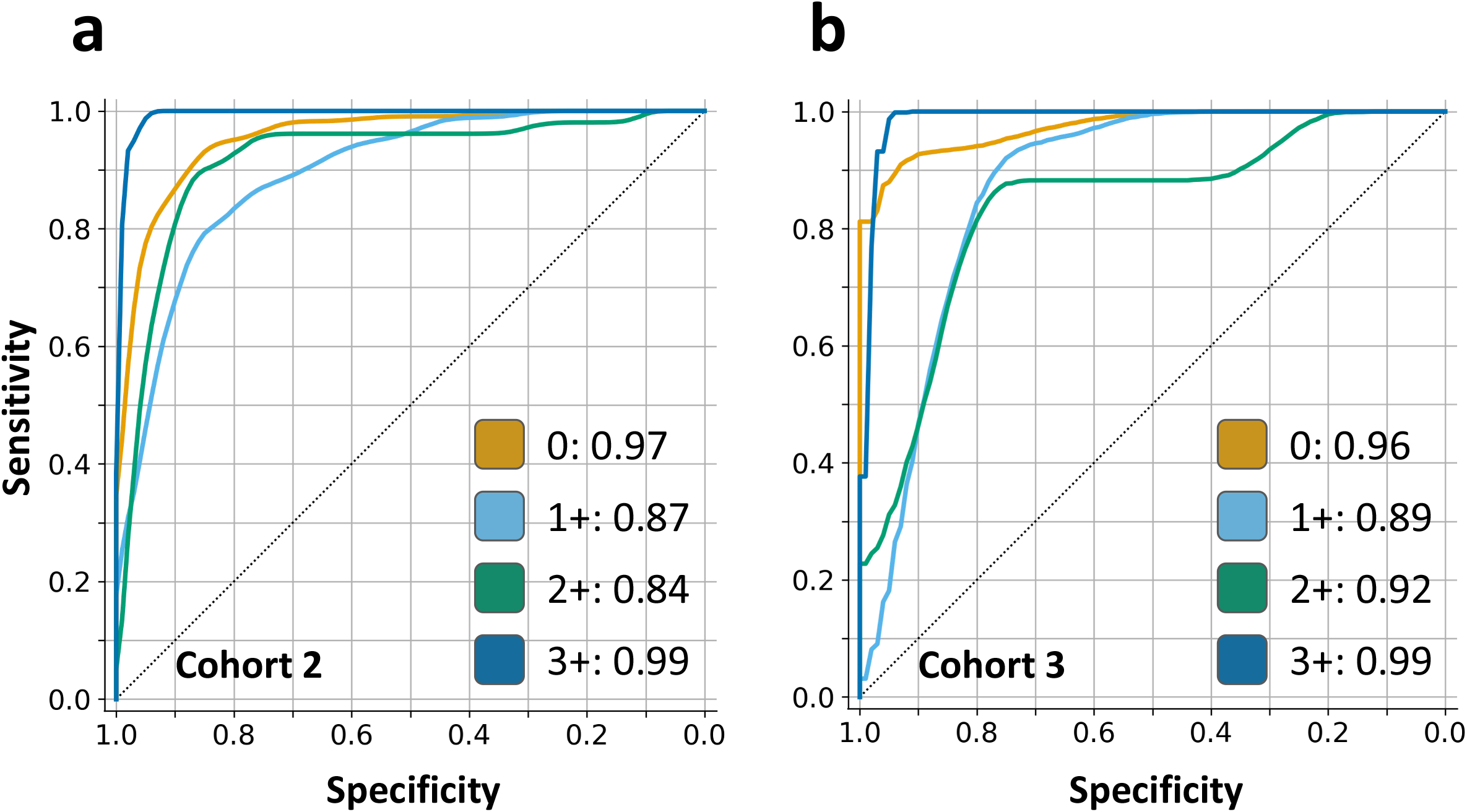
External validation of the HER2 IHC model. a) ROC curves of the AI IHC model on Cohort 2. Each curve is a one-vs-rest assessment of the model’s ability to distinguish every IHC score from the other. **b)** ROC curves of the AI IHC model on Cohort 3. The most difficult subtask consists of distinguishing between IHC 1+ (HER2-low) and IHC 0 (HER2-negative) cases.

### The model identifies HER2-low tumors with high sensitivity

The model excelled in differentiating between HER2-negative (0) and non-HER2-negative (1+, 2+, 3+) tumors. The model was very sensitive to HER2-low and HER2-positive tumors of 0.95 [0.93 - 0.98] on Cohort 2 and 0.97 [0.91 - 1.00] on Cohort 3. Moreover, the model maintained specificity for non-HER2-negative tumors, with specificity scores of 0.79 [0.72 - 0.84] for Cohort 2 and 0.72 [0.63 - 0.82] for Cohort 3.

We did not find a significant change in performance on core biopsies (Cohort 2, AUC = 0.95 [0.93 = 0.97], Sensitivity 0 vs 1+/2+/3+ = 0.95 [0.92 - 0.98]) and surgical resections (Cohort 2, AUC = 0.94 [0.91 - 0.97], Sensitivity 0 vs 1+/2+/3+ = 1.00 [0.98 - 1.00]). Besides, we found no change in performance with respect to the hormone receptor status (**Supplementary Table 4**).

To validate the relevance of the features our model captures for IHC assessment, we conducted an ablation study using features from a model pre-trained only on H&E tiles (**Supplementary Table 5**). The resulting AUC of 0.92 [0.90 - 0.94] was significantly lower than that of our model pre-trained on IHC (DeLong P < 0.001), affirming that the features learned by our model are significant for evaluating HER2 protein levels.

#### The IHC model to assess HER2 expression levels mirrors the ASCO/CAP guidelines

To ensure our model accurately quantified HER2 from IHC slides, we presented the highest and lowest-scoring predictive tiles to a pathologist for verification (**Figure 3a**). The IHC model captures the completeness of the membrane staining, as seen in the top predictive tiles across all slides of Cohort 2 (**Figure 3b**). The top predictive tiles for scores 1+ and 2+ closely adhere to the staining patterns described in the ASCO/CAP guidelines for HER2 testing. The predictive tiles for the 1+ score demonstrate weak and partial membrane staining. In contrast, the tiles for the 2+ score show a heterogeneous expression pattern with a combination of a weak complete membrane and a moderate incomplete membrane staining.

**Figure 3:**
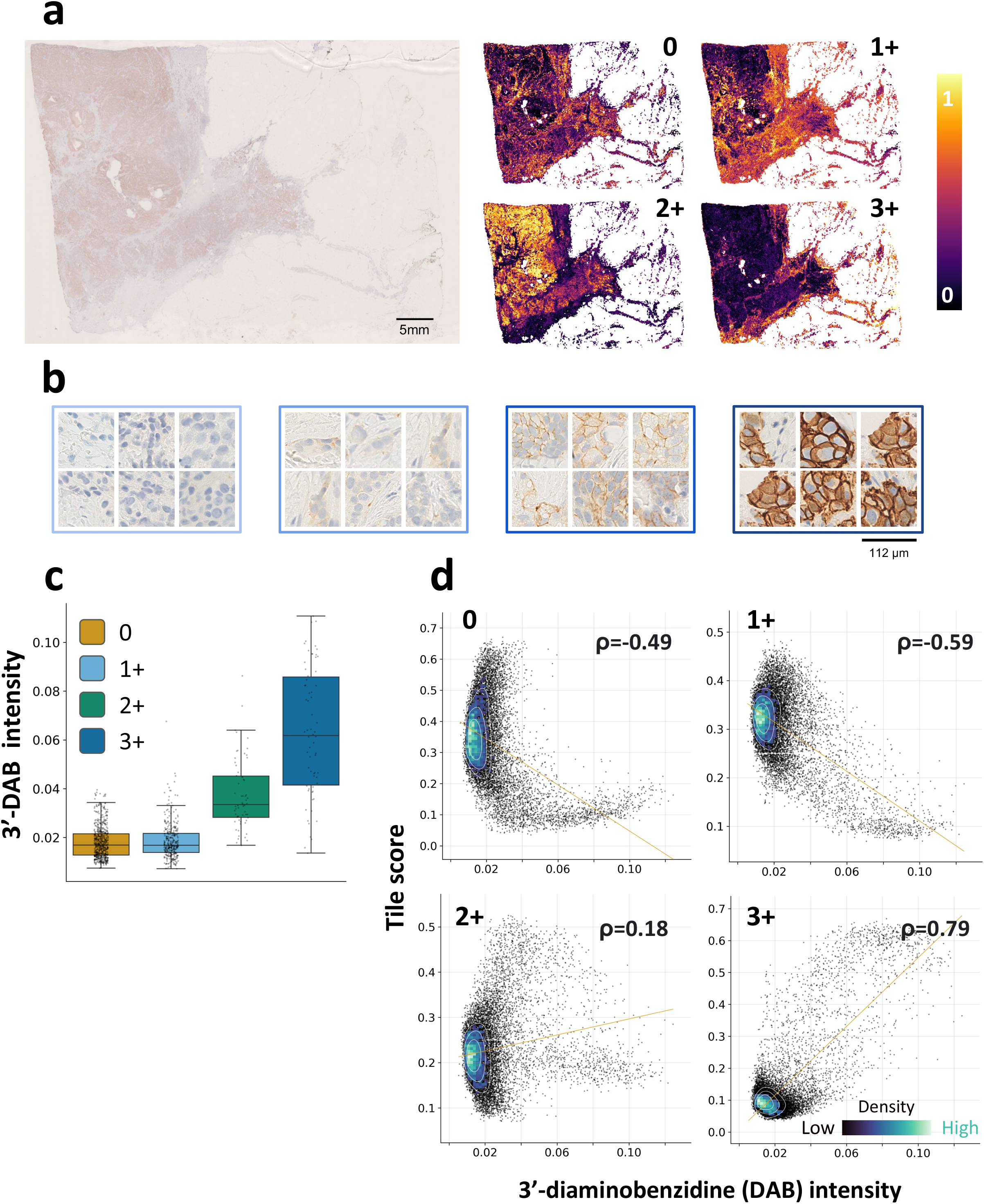
The AI IHC model mirrors the 2018 ASCO/CAP guidelines. a) The AI IHC model prediction heatmaps showing a probability of being 0 / 1+ / 2+ / 3+ for each 112 x 112 μm tile of the slide. **b)** Most predictive tiles across all slides of Cohort 2 for each IHC score category; from left to right 0, 1+, 2+ and 3+. The first image (0) shows no staining at all. The second image (1+) shows weak and partial membrane staining. The third image (2+) shows a weak complete membrane to moderate incomplete membrane staining. The AI IHC model not only captures how strong the membrane staining is but also the completeness of the membrane staining, a key differentiator of IHC 1+ and IHC 2+ tiles. Finally, the last image (3+) shows strong complete membrane staining. These predictive tiles closely adhere to the staining patterns from the ASCO-CAP 2018/2023 guidelines for HER2 testing. **c)** Distribution of the DAB channel intensity per IHC score category. Each grey dot represents a tile. The IHC score assigned to the tile corresponds to the IHC score of the slide predicted by the model. **d)** Correlation of the 3’- diaminobenzidine (DAB) channel intensity and the probability of being 0 / 1+ / 2+ / 3+ attributed by the AI model to each 112 x 112 μm tile of the slides from Cohorts 2 and 3.

To validate that the model was assessing HER2 levels by applying the ASCO/CAP guidelines, we investigated the correlation between the scores output by the model to the tiles and the amount of 3’-diaminobenzidine (DAB) on the tiles. For each slide of Cohort 2 and 3, we selected the 100 highest-scoring tiles for all the IHC score categories (0, 1+, 2+, and 3+) and extracted their DAB channel. The DAB intensity correlated with the IHC scores given by the model (Kruskal-Wallis P < 0.001, **Figure 3c**). The correlation between the model scores and the intensity of the DAB channel (**Figure 3d**) was significant for the IHC 0 category (ρ=-0.49, Fisher P < 0.001), 1+ (ρ=-0.59, Fisher P < 0.001) and 3+ (ρ=0.79, Fisher P < 0.001) categories. While identified by our model, the tiles considered predictive of the IHC 2+ score did not significantly correlate (ρ=0.18) with the DAB channel intensity.

#### Ambiguous cases have clear morphological features and artifacts

To characterize the histomorphological features associated with the different levels of HER2 expression, we selected 25 pairs of H&E and IHC slides from Cohort 2 and registered them. We computed the correlation coefficients for each of the HER2-positive, HER2-low, and HER2-negative heatmaps for every pair of registered slides (**Figure 4a, 4b**). For the HER2- positive (respectively HER2-low and HER2-negative) heatmaps, we found 9 (respectively 13 and 10) slides with a statistically significant positive correlation between the H&E and IHC heatmaps with an average Pearson correlation of 0.22 [0.14 - 0.31] (respectively 0.20 [0.14 - 0.27] and 0.24 [0.15 - 0.33]). We ran an in-house histomics model trained on TCGA to predict 5 tissue types: tumor, stroma, lymphocytes, fat, and necrosis (**Figure 4c**). The correlation analysis showed that the HER2-positive status was mainly found in tiles exhibiting tumor cells, while it was negatively associated with the presence of necrosis and lymphocytes (**Figure 4d**). The tiles predictive of the HER2-low and HER2-negative statuses exhibit the same tissue content, hinting at the difficulty of distinguishing the HER2-low and the HER2-negative statuses directly on H&E slides. These tiles showed mostly stroma content and, more precisely, regions exhibiting fibrosis. On the opposite side, the HER2-negative status is associated with the presence of lymphocytes and necrosis (**Figure 4d**).

**Figure 4:**
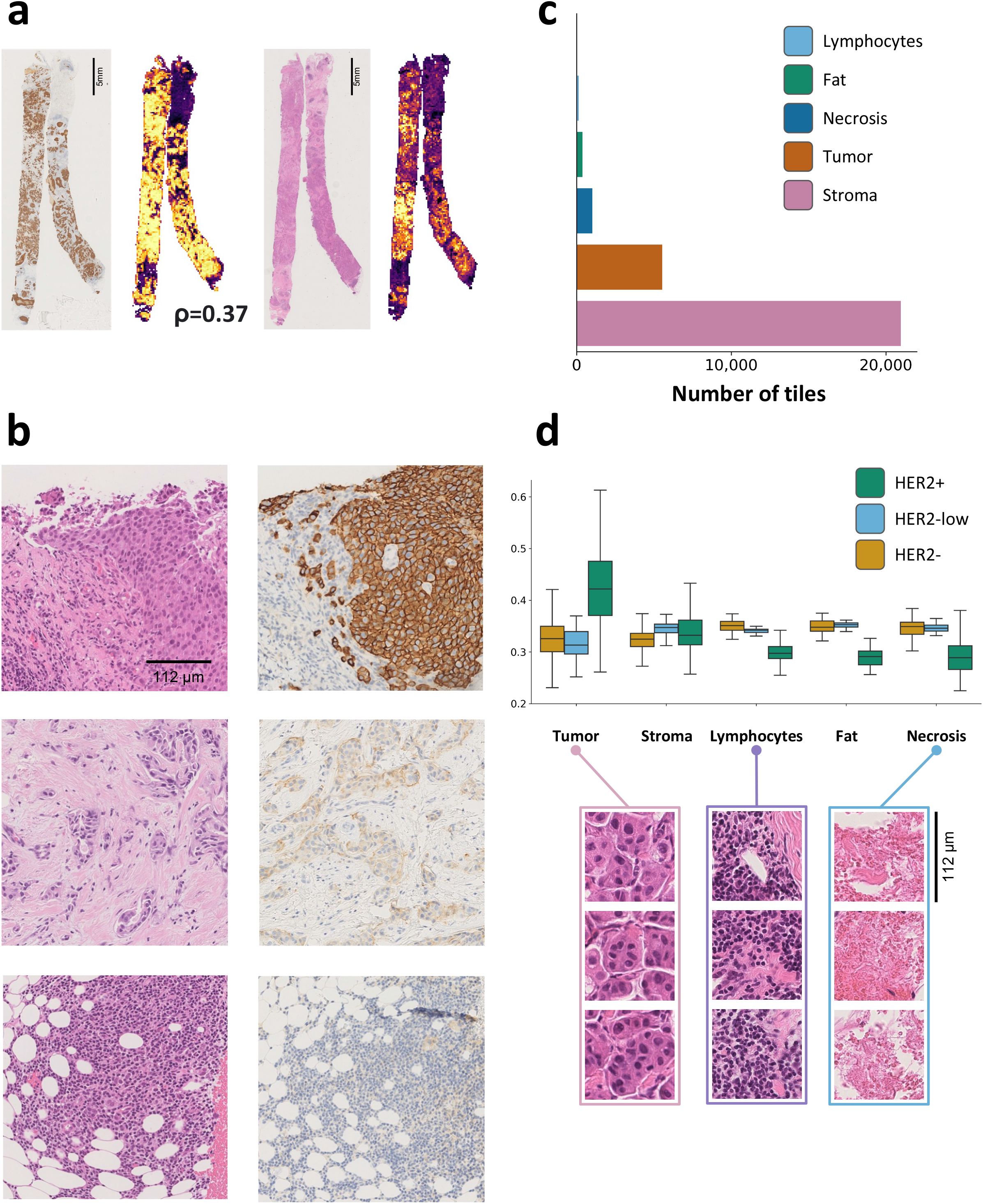
Tumor heterogeneity based on the HER2 status. a) Pairs of H&E and IHC slides registered with their corresponding heatmaps of scores assigned by the AI IHC and H&E model. On the left, both heatmaps correspond to the probability of a tile being HER2+. On the right, the heatmaps exhibit the probability of being HER2-low. The number on the bottom is the correlation coefficient between scores assigned by the IHC and H&E model to the same tile. **b)** Matching H&E and IHC regions predictive of the HER2-positive status (top), HER2-low status (middle), HER2-negative status (bottom). These matching regions were obtained by registering pairs of consecutive H&E and IHC slides, and found by looking at the most predictive regions of the H&E and IHC models for the 3 HER2 statuses. **c)** Histomics distribution on H&E slide of figure a). **d)** Distribution of histological patterns with respect to the HER2 status.

Finally, we analyzed the artifacts that prevented the DL model from accurately distinguishing HER2-null from HER2-low. These artifacts included glue residue under the coverslip (cases 1 and 2 in **Supplementary** Figure 1), non-specific cytoplasmic (cases 1, 5, and 6 in **Supplementary** Figure 1 and 2), stromal, and artefactual fibrin or blood staining (cases 7, 8 and 9 in **Supplementary** Figure 2 and 3). Additionally, cases with minimal HER2 expression (1+) among normal glands (case 3 in **Supplementary** Figure 1) complicated classification. Physical damage, such as crushed tissue (case 4 in **Supplementary** Figure 1) and fixation artifacts, made the scoring less accurate, leading to potential misclassification by the DL model.

## Discussion

The emergence of anti-HER2 antibody-drug conjugates (ADCs) as a therapeutic option for HER2-low breast cancer underscores the critical need for precise HER2 status assessment. In this study, we developed a deep learning (DL) model to complement the ASCO-CAP criteria, aiding pathologists in interpreting HER2-Low cases. By evaluating a comprehensive, multi-centric dataset comprising 760 cases, our results indicate that the DL model proficiently identifies up to 97% of HER2-low and HER2-positive cases. This capability holds significant clinical relevance as it identifies patients who may benefit from therapies such as trastuzumab deruxtecan (T-DXd), thereby underscoring the practical applicability of our model.

Our study demonstrates that the DL model enhances concordance among pathologists while also identifying key factors contributing to discordance between pathologists and the DL model. Notably, factors such as extensive fibrosis, high tumor-infiltrating lymphocyte counts, and necrosis, as well as common artifacts in immunohistochemistry, like non-specific background cytoplasmic staining, contribute to such discrepancies.

Our developed models are designed for interpretability, assigning scores to individual tiles within whole-slide images and mirroring ASCO/CAP scoring guidelines. The features extracted from the tiles detect the right information on different intensities of membrane staining and their location on the cells. The model subsequently correctly maps these to the appropriate IHC score. For the IHC 1+ and 2+ categories, we show that the models’ scores were less explained by the intensity of staining on the tiles and more by the completeness of the cell membrane staining.

Moreover, our study further highlights the spatial heterogeneity of HER2 expression. By analyzing 25 consecutive slides of breast carcinomas, one for the H&E slide and one for the IHC, we were able to meticulously identify areas predictive of the HER2-negative, HER2-low, and HER2-positive status, offering deeper insights into the morphological characteristics of HER2-expression in these areas.

The predictive tiles for HER2-positive cases are clearly characterized by tumoural cellular and nuclear details. In contrast, the top predictive tiles for the ambiguous HER2-low cases were mostly non-specific features in the stroma, such as blood for example. The non-specific properties of these top predictive tiles might make distinguishing HER2-negative from HER2- low cases difficult. Furthermore, specific and well-known artifacts of immunohistochemistry, such as background staining in the cytoplasm of tumor cells and stroma, can make the DL model unable to differentiate between cases with no expression at all (HER2-null) versus some expression (HER2-ultralow). This is important as it illustrates that only by co-use of a DL model, along with the expertise of the pathologist, with knowledge of the artifacts and morphological features described in this study, the odds of misclassifying a HER2-low case can be limited.

Given the clinical significance of antibody-drug conjugates such as T-Dxd for breast cancer patients^2^, precise identification of HER2-low cases is imperative. Although we have not extensively explored HER2-ultralow versus HER2-null assessment using our method, future considerations may arise pending the Destiny06 Trial outcomes, potentially necessitating cohort re-annotation to accommodate a HER2-ultra-low category^7^.

## Materials and methods

### Datasets description

The training set, “Cohort 1”, is comprised of 299 cases from 16 centers in région Rhône-Alpes-Auvergne (2018-2020) with IHC slides (HER2/neu 4B5 Ventana assay), scanned with an Aperio Leica GT 450 at 40X. Samples for the validation cohort, “Cohort 2”, were obtained from The King’s Health Partners (KHP) Cancer Biobank at Guy’s Hospital (London, UK; REC no.: 07/40874/131). Basic clinical characteristics for this cohort were available and shown in **Supplementary Table 6**. Cohort 3, an independent Polish cohort, includes 92 cases. Both cohorts have slides for HER2 immunohistochemistry (HER2/neu 4B5 Ventana assay) and H&E, scanned using Hamamatsu scanners at 40X magnification.

### Annotation of HER2 immunohistochemistry slides

The three cohorts, Cohort 1, Cohort 2, and Cohort 3, were annotated using the ASCO/CAP 2018/2023 guidelines without access to any metadata. Cohort 1 and Cohort 3 were each annotated by one expert pathologist. Cohort 2 was independently annotated by 3 breast expert pathologists, with the ground truth defined as a majority vote between the three annotators.

#### Preprocessing of whole-slide images

The preprocessing of WSI can be decomposed into 3 steps: 1) matter detection, 2) tiling, and 3) feature extraction. Using a U-Net neural network, tissue on WSIs is identified and segmented, discarding irrelevant areas. This network was trained on 460 H&E and IHC slides, achieving a Dice score of 0.96 in validation. Next, the identified tissue regions are divided into 224x224 pixel tiles (effective area of 112x112 micrometers). Tiles from H&E slides require at least 60% tissue presence, whereas IHC slides only need 10% to ensure comprehensive coverage. Then, 2048 features are extracted for each tile using a wide ResNet50 network^8^. The feature extractors for H&E and IHC slides are trained in a self-supervised fashion using the approach described in ^9^ on 4 million tiles from the TCGA-COAD dataset with massive data augmentation and without using any labels. The feature extractor for IHC slides is trained on 2 million tiles from the HER2 Cohort 1 dataset using the same self-supervised framework.

#### HER2 status prediction

For training and inference, 5,000 tiles for IHC and H&E are uniformly sampled from each slide for speed and memory considerations. Both models have an identical structure, as proposed in ^9^. The output scores are passed to a softmax layer to obtain probabilities. The models were trained with the categorical cross entropy as a loss function, using the labels obtained from pathologists as described in the section “Annotation of HER2 immunohistochemistry slides”.

#### Performance assessment and statistical methods

The area under the receiver operating characteristic (AUC) one-vs-one (OVO) was used to quantify the capability of the model to distinguish the 4 IHC scores. The sensitivity and specificity of HER2-low and HER2-positive were used to monitor the ability of the model to identify HER2-low cases. Confidence intervals at 95% confidence level were obtained by bootstrapping experiment results with 1000 repeats. Pearson’s correlation was used to quantify the correlation between the H&E and IHC tile scores. All tests were two-tailed, and P-values < 0.05 were considered statistically significant. The intraclass-correlation coefficient (ICC) was computed using the Pingouin library and corresponds to the ICC2k (Average random raters).

#### Slide registration

To validate the regions picked by the H&E model as highly predictive of the HER2 status, 25 pairs of H&E and IHC slides from Cohort 2 were selected. The slides were registered by finding the optimal Euler (rotation) and Spline (scaling and shearing) transforms by maximizing the mattes advanced mutual information criterion via gradient descent on ten resolution levels. The registration was carried out using the Elastix library and validated by visual inspection to ensure the registered tiles matched.

## Acknowledgments

We thank Marie Brevet for providing Cohort 1.

## Author contributions

Study conception and design: S.I, R.S., C.M. and P.-A.B.; Data collection: L.V.P., E.A., A.M., J.T., C.G., E.S., P.Z.; Data annotation: G.B., R.S., P.G., M.L.T.; Software: P.-A.B., L.H., R.D.; Analysis and interpretation of results: P.-A.B., G.B., F.D., S.I., R.S.; Draft manuscript preparation: P.-A.B., G.B, R.S. S.I. All authors reviewed the results and approved the final version of the manuscript.

---

**Supplementary Table 1:** Distribution of slide artifacts for the validation cohorts.

| Artefacts | Number of instances |  |
| --- | --- | --- |
|  | Validation Cohort 2 | Validation Cohort 3 |
| Out-of-focus<sup>(1)</sup> | 5 | 0 |
| Pen markers | 14 | 0 |
| Little to no tumor | 12 | 0 |
| Fixation issues | 9 | 0 |
| Air bubbles | 2 | 0 |
| Folded tissue | 2 | 5 |
| Low-quality staining<sup>(2)</sup> | 7 | 9 |
| Miscellaneous<sup>(3)</sup> | 8 | 4 |

<sup>(1)</sup> Slides with more than 50% of out-of-focus regions rendering the HER2 IHC score assessment impossible
<sup>(2)</sup> Cytoplasmic or nuclei staining on part or the full slide
<sup>(3)</sup> Low-quality samples pathologists could not assess

**Supplementary Table 2:** Area under the operating characteristic curve (AUROC) per class (0, 1+, 2+, 3+) of the IHC model on 2 external validation cohorts.

| Cohorts | OVO AUC | OVR AUC |  |  |  |
| --- | --- | --- | --- | --- | --- |
|  |  | 0 | 1+ | 2+ | 3+ |
| **Validation Cohort 2** | 0.95<br>[0.93 - 0.96] | 0.95<br>[0.94 - 0.95] | 0.80<br>[0.79 - 0.82] | 0.91<br>[0.90 - 0.92] | 0.99<br>[0.99 - 1.00] |
| **Validation Cohort 3** | 0.92<br>[0.90 - 0.93] | 0.97<br>[0.96 - 0.98] | 0.78<br>[0.76 - 0.81] | 0.84<br>[0.80 - 0.88] | N/A |

**Supplementary Table 3:** Confusion matrices on Cohort 2 (a) and Cohort 3 (b) for the IHC model outputting a HER2 IHC score.

**a**
| Ground truth | Predicted |  |  |  |  |
| --- | --- | --- | --- | --- | --- |
|  |  | 0 | 1+ | 2+ | 3+ |
|  | 0 | 91 | 26 | 1 | 4 |
|  | 1+ | 8 | 72 | 14 | 1 |
|  | 2+ | 0 | 4 | 40 | 12 |
|  | 3+ | 0 | 0 | 1 | 50 |

**b**
| Ground truth | Predicted |  |  |  |  |
| --- | --- | --- | --- | --- | --- |
|  |  | 0 | 1+ | 2+ | 3+ |
|  | 0 | 39 | 4 | 0 | 0 |
|  | 1+ | 1 | 20 | 0 | 0 |
|  | 2+ | 0 | 7 | 1 | 1 |
|  | 3+ | 0 | 0 | 0 | 1 |

**Supplementary Table 4:** Comparison of performance of the IHC model with respect to the hormone receptor status on Cohort 2 for estrogen receptor (a) and progesterone receptor (b).

**a**
| Validation Cohort 2 | AUC OVO | Sensitivity (0 vs 1+/2+/3+) | Specificity (0 vs 1+/2+/3+) |
| --- | --- | --- | --- |
| ER+ | 0.97<br>[0.96 - 0.98] | 0.95<br>[0.93 - 0.97] | 0.68<br>[0.62 - 0.75] |
| ER- | 0.97<br>[0.96 - 0.98] | 0.97<br>[0.93 - 1.00] | 0.73<br>[0.62 - 0.83] |

**b**
| Validation Cohort 2 | AUC OVO | Sensitivity (0 vs 1+/2+/3+) | Specificity (0 vs 1+/2+/3+) |
| --- | --- | --- | --- |
| PR+ | 0.94<br>[0.92 - 0.95] | 0.96<br>[0.93 - 0.98] | 0.66<br>[0.58 - 0.75] |
| PR- | 0.96<br>[0.94 - 0.97] | 0.95<br>[0.91 - 0.98] | 0.77<br>[0.70 - 0.84] |

**Supplementary Table 5:** Ablation study comparing two HER2 IHC models (feature extractors): one is pre-trained only on H&E tiles, the other only on IHC tiles.

| Validation Cohort 1 | AUC OVO | Sensitivity (0 vs 1+/2+/3+) | Specificity (0 vs 1+/2+/3+) |
| --- | --- | --- | --- |
| Pre-training on IHC tiles | 0.95<br>[0.93 - 0.96] | 0.95<br>[0.93 - 0.98] | 0.79<br>[0.72 - 0.84] |
| Pre-training on H&E tiles | 0.92<br>[0.90 - 0.94] | 0.97<br>[0.94 - 0.99] | 0.53<br>[0.45 - 0.59] |

**Supplementary Table 6:** Clinical variables for Cohort 2. We present descriptive statistics (mean, median, min-max values) for the continuous variables. The figure in parenthesis correspond to the standard deviations. For categorical variables, the absolute frequencies (as well as the relative frequencies in parentheses) are presented. The “Unknown” category signifies that we could not obtain the information about some patients.

| Variables |  | Cohort 2 (n=369) |
| --- | --- | --- |
| Age (years) | Mean | 54.5 (14.5) |
|  | Median | 52.4 |
|  | Min-Max | 22-94 |
| Year of diagnosis | Mean | 2018 |
|  | Median | 2018 |
|  | Min-Max | 1985 - 2021 |
| Ethnic group | White | 193 (52.3%) |
|  | Black | 55 (14.9%) |
|  | Asian | 14 (3.8%) |
|  | Mixed | 6 (1.7%) |
|  | Other | 13 (3.5%) |
|  | Unknown | 88 (23.8%) |
| ER Status | Negative | 70 (19.0%) |
|  | Positive | 271 (73.4%) |
|  | Unknown | 28 (7.6%) |
| PR Status | Negative | 118 (32.0%) |
|  | Positive | 215 (58.3%) |
|  | Unknown | 36 (9.7%) |
| In-situ type | DCIS | 208 (56.4%) |
|  | LCIS | 28 (7.6%) |
|  | Mixed | 5 (1.4%) |
|  | Unknown | 128 (34.6%) |
| Grade | I | 25 (6.8%) |
|  | II | 102 (27.6%) |
|  | III | 79 (21.4%) |
|  | Unknown | 163 (44.2%) |

**Supplementary Figure 1:**
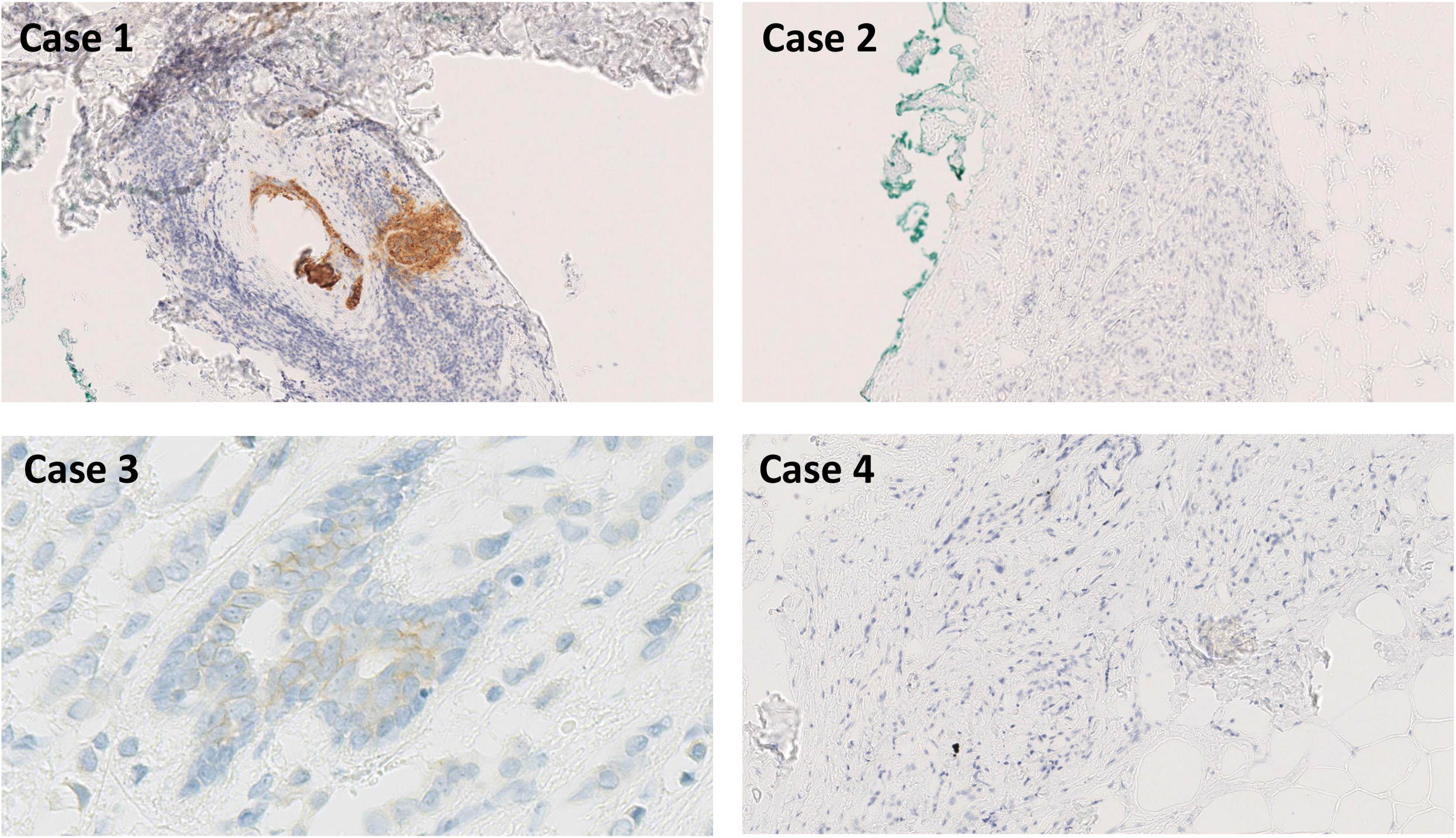
A collection of artefacts preventing an accurate distinction between HER2-null and HER2-ultra-low cases. Case 1 and 2 present glue under the coverslip. Case 1 also presents a 3+ DCIS foci. Case 3 presents normal glands stain 1+. Case 4 presents crushed invasive cancer cells.

**Supplementary Figure 2:**
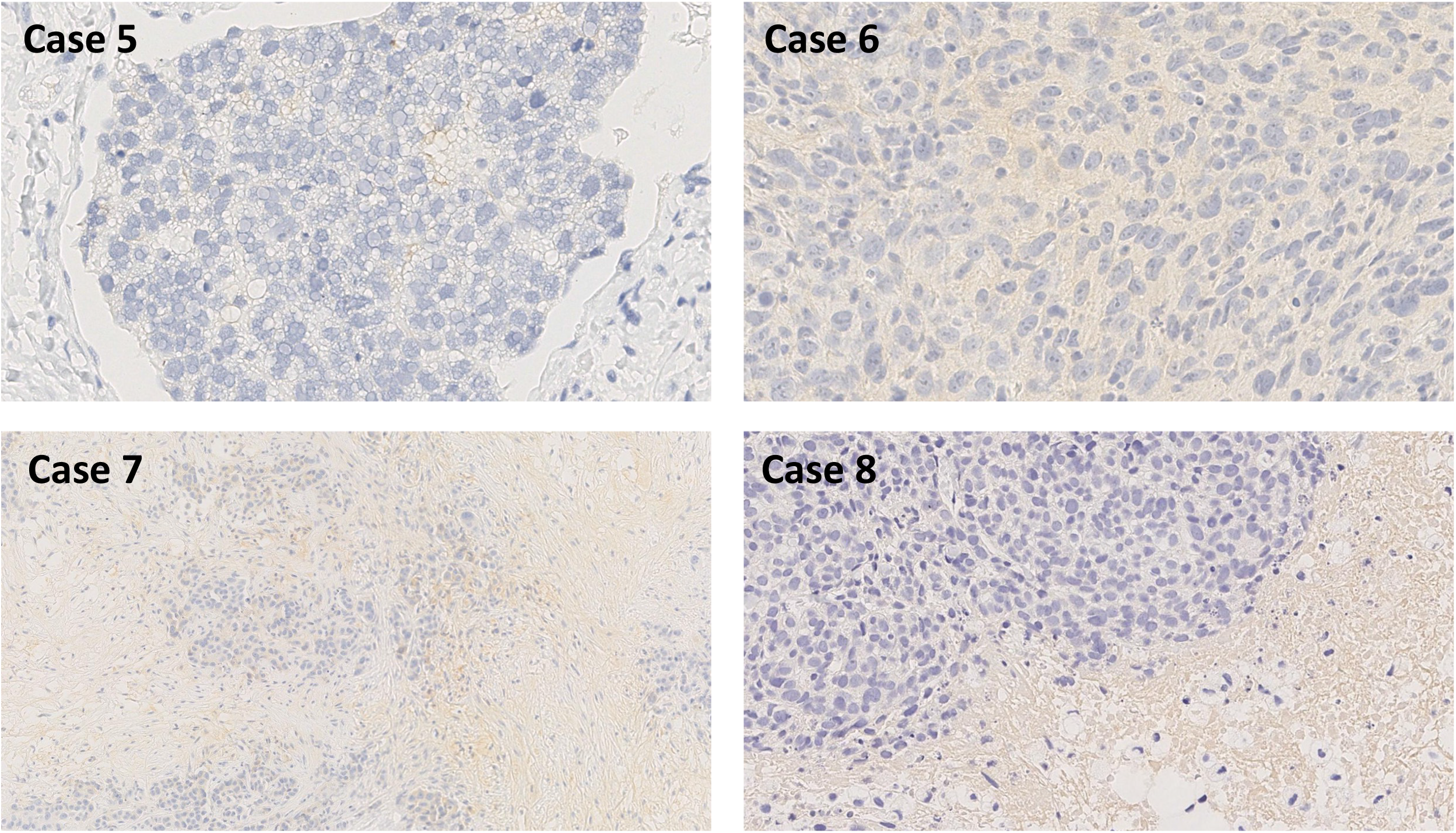
Case 5 and 6 present faint cytoplasmic stain in tumor cells and stroma. Case 7 and 8 present non-specific stain in fibrin and blood.

**Supplementary Figure 3:**
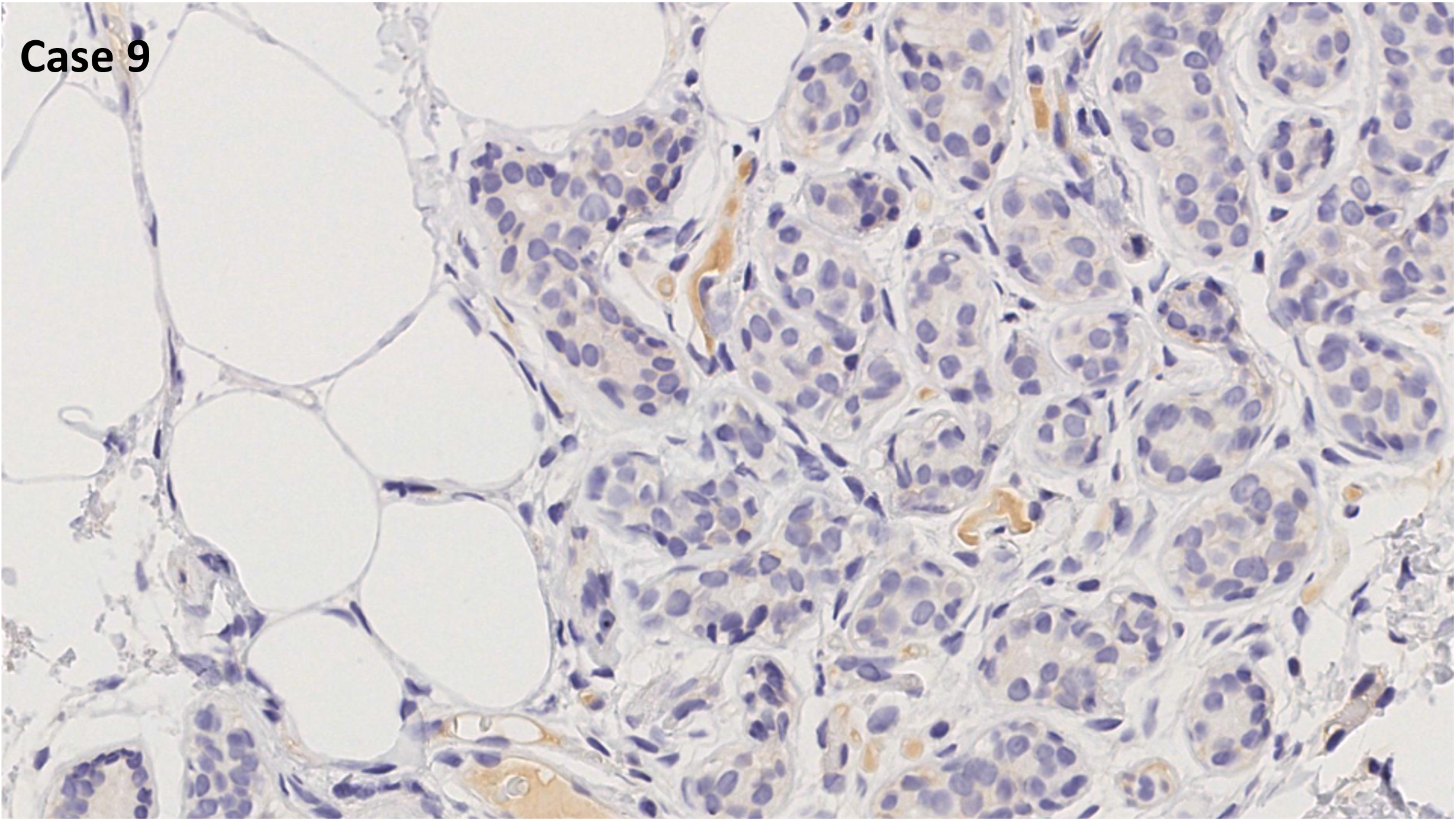
Case 9 present artifactual fibrin/blood stain.

